# *In vivo* imaging of cannabinoid type 2 receptors, functional and structural alterations in mouse model of cerebral ischemia by PET and MRI

**DOI:** 10.1101/2021.05.08.441033

**Authors:** Ruiqing Ni, Adrienne Müller Herde, Ahmed Haider, Claudia Keller, Georgios Louloudis, Markus Vaas, Roger Schibli, Simon M. Ametamey, Jan Klohs, Linjing Mu

## Abstract

**Background and purpose:** Brain ischemia is one of the most important pathologies of the central nervous system. Non-invasive molecular imaging methods have the potential to provide critical insights into the temporal dynamics and follow alterations of receptor expression and metabolism in ischemic stroke. The aim of this study was to assess the cannabinoid type 2 receptors (CB_2_R) levels in transient middle cerebral artery occlusion (tMCAO) mouse models at subacute stage using positron emission tomography (PET) with our novel tracer [^18^F]RoSMA-18-d6, and structural imaging by magnetic resonance imaging (MRI).

**Methods:** Our recently developed CB_2_R PET tracer [^18^F]RoSMA-18-d6 was used for imaging the neuroinflammation at 24 h after reperfusion in tMCAO mice. The RNA expression levels of CB_2_R and other inflammatory markers were analyzed by quantitative real-time polymerase chain reaction using brain tissues from tMCAO (1 h occlusion) and sham-operated mice. [^18^F]fluorodeoxyglucose (FDG) was included for evaluation of the cerebral metabolic rate of glucose (CMRglc). In addition, diffusion-weighted imaging and T_2_-weighted imaging were performed for anatomical reference and delineating the lesion in tMCAO mice.

**Results:** mRNA expressions of inflammatory markers *TNF-α*, *Iba1, MMP9* and *GFAP, CNR2* were increased at 24 h after reperfusion in the ipsilateral compared to contralateral hemisphere of tMCAO mice, while mRNA expression of the neuronal marker *MAP-2* was markedly reduced. Reduced [^18^F]FDG uptake was observed in the ischemic striatum of tMCAO mouse brain at 24 h after reperfusion. Although higher activity of [^18^F]RoSMA-18-d6 in *ex-vivo* biodistribution studies and higher standard uptake value ratio (SUVR) were detected in the ischemic ipsilateral compared to contralateral striatum in tMCAO mice, the *in-vivo* specificity of [^18^F]RoSMA-18-d6 was confirmed only in the CB_2_R-rich spleen.

**Conclusions:** This study revealed an increased [^18^F]RoSMA-18-d6 measure of CB_2_R and a reduced [^18^F]FDG measure of CMRglc in ischemic striatum of tMCAO mice at subacute stage. [^18^F]RoSMA-18-d6 might be a promising PET tracer for detecting CB_2_R alterations in animal models of neuroinflammation without neuronal loss.

## Introduction

The pathophysiology of ischemic stroke is complex and associated with a myriad of cellular and molecular pathways. The severe reduction in cerebral blood flow (CBF) initiates a cascade of hemodynamic, vascular and inflammatory processes in a time-dependent manner in the supplied brain territory, and subsequent defensive response for repair related to lesion expansion and containment. Irreversible tissue damage occurs in the core of the ischemic area; while neurons in the ischemic penumbra face excitotoxicity, peri-infarct polarizations, inflammation and apoptosis, leading to a secondary tissue damage and expansion of the lesion if reperfusion cannot be restored within an early time frame [2–4]. Neuroinflammation post stroke has been an important therapeutic target. Anti-inflammatory, immunomodulatory treatments and microglia-targeted therapy were evaluated in clinical stroke trials [5–7]. Thus, there is a need for imaging the regional neuroinflammatory pattern for understanding disease mechanism and for therapeutic monitoring.

Positron emission tomography (PET) using [^18^F]fluorodeoxyglucose ([^18^F]FDG) for cerebral metabolic rate of glucose (CMRglc), [^15^O]H_2_O for perfusion imaging, and diffusion weighted (DW) magnetic resonance imaging (MRI) are valuable tools to support understanding of the pathophysiology in patients with ischemic stroke [3, 8–14]. However, *in vivo* imaging of neuroinflammation and gliosis is challenging [12, 13, 15]. One reason is that the astrocytes and microglia are highly dynamic and heterogeneous in their subtypes, locations and activation status. Additionally, the identification of an ideal target for neuroinflammation imaging is highly demanding. Translocator protein (TSPO) is the most widely used neuroinflammation target for PET imaging. [^11^C]PK-11195, the first generation TSPO PET tracer, and several second-generation tracers such as [^11^C]DAA1106, [^11^C]PBR06, [^11^C]PBR28, [^11^C]GE180, and [^18^F]DPA-713, [^18^F]DPA-714 [16–24] have been evaluated in (pre)-clinical studies. So far, imaging neuroinflammation with TSPO PET tracers yielded controversial results in rodents and patients with ischemic stroke [1, 13, 20]. Thus, the development of novel PET probes for visualizing alternative targets in neuroinflammation have received great attention in recent years [25–27].

Cannabinoid type 2 receptors (CB_2_R) are mainly expressed by immune cells including monocytes and macrophages. In the brain, CB_2_Rs are primarily found on microglia and have low expression levels under physiological conditions [2, 4, 28]. Upregulation of brain CB_2_R expression is reported under acute inflammation such as ischemic stroke, and related to lesion extension in the penumbra and subsequent functional recovery [29]. Treatment with CB_2_R agonists has been shown to be neuroprotective and attenuates macrophage/microglial activation in the mouse models of cerebral ischemia [29, 42–45]. CB_2_R is also upregulated in other brain diseases with involvement of inflammation/microglia under chronic inflammation in neurodegenerative diseases such as Alzheimer’s disease [30–33] and senescence-accelerated models [34], associated with amyloid-β deposits[28, 35–41].

Several structural scaffolds of CB_2_R PET tracers have recently been developed [46–50] including pyridine derivatives, oxoquinoline derivatives; thiazole derivatives [51, 52]; oxadiazole derivatives [53]; carbazole derivatives [54]; imidazole derivative [55]; and thiophene derivatives [56, 57]. In this study, our newly developed pyridine derivative [^18^F]RoSMA-18-d6, which exhibited sub-nanomolar affinity and high selectivity towards CB2R (Ki: 0.8 nM, CB2R/CB1R > 12’000) [58] is selected as the CB_2_R PET tracer.

The aim of the current study was to evaluate the novel CB_2_R tracer [^18^F]RoSMA-18-d6 in the transient middle cerebral artery occlusion (tMCAO) mouse models of focal cerebral ischemia [60–66] using microPET. In addition, [^18^F]FDG was included for evaluation of the cerebral metabolic rate of glucose (CMRglc). Diffusion-weighted imaging and T_2_- weighted imaging were performed for anatomical reference and for delineating the lesion in tMCAO mice.

## Methods

### Radiosynthesis

[^18^F]RoSMA-18-d6 was synthesized by nucleophilic substitution of the tosylate precursor with [^18^F]KF/Kryptofix222 in acetonitrile [58]. The crude product was purified by reverse phase semi-preparative high-performance liquid chromatography and formulated with 5 % ethanol in water for intravenous injection and for biological evaluations. In a typical experiment, a moderate radiochemical yield of ~ 12 % (decay corrected) was achieved with a radiochemical purity > 99 %. The molar activities ranged from 156 to 194 GBq/μmol at the end of synthesis. The identity of the final product was confirmed by comparison with the HPLC retention time of the non-radioactive reference compound by co-injection. [^18^F]FDG was obtained from a routine clinical production from the University Hospital Zurich, Switzerland.

### Animals

Twenty-four male C57BL/6J mice were obtained from Janvier Labs (Le Genest-Saint-Isle, France). The mice were scanned at 8–10 weeks of age (20–25 g body weight). Mice were randomly allocated to sham-operation (n = 10) or tMCAO (n = 14). Mice underwent MRI, μPET/ computed tomography (CT), and 2,3,5-Triphenyltetrazolium chloride (TTC) histology staining for validation 24 h or 48 h after reperfusion. Animals were housed in ventilated cages inside a temperature-controlled room, under a 12-hour dark/light cycle. Pelleted food (3437PXL15, CARGILL) and water were provided *ad-libitum*. Paper tissue and red Tecniplast mouse house® (Tecniplast, Milan, Italy) shelters were placed in cages as environmental enrichments. All experiments were performed in accordance with the Swiss Federal Act on Animal Protection and were approved by the Cantonal Veterinary Office Zurich (permit number: ZH018/14 and ZH264/16).

Surgeries for tMCAO and sham-operation were performed using standard-operating procedures as described before [67, 68]. Anaesthesia was initiated by using 3 % isoflurane (Abbott, Cham, Switzerland) in a 1:4 oxygen/air mixture, and maintained at 2 %. Before the surgical procedure, a local analgesic (Lidocaine, 0.5 %, 7 mg/kg, Sintectica S.A., Switzerland) was administered subcutaneously (s.c.). Temperature was kept constant at 36.5 ± 0.5 °C with a feedback controlled warming pad system. All surgical procedures were performed in 15-30 min. After surgery, buprenorphine was administered as s.c. injection (Temgesic, 0.1 mg/kg b.w.), and at 4 h after reperfusion and supplied thereafter via drinking water (1 mL/32 mL of drinking water) until 24 h or 48 h. Animals received softened chow in a weighing boat on the cage floor to encourage eating. tMCAO animals were excluded from the study if they met one of the following criteria: Bederson testing was performed 2h post-reperfusion. Bederson score of 0, no reflow after filament removal, and premature death.

#### mRNA isolation, reverse-transcription reaction and real-time polymerase chain reaction

Brain hemispheres of C57BL/6 mouse, tMCAO mice at 24 h and 48 h post reperfusion were used for total mRNA isolation according to the protocols of the Isol-RNA Lysis Reagent (5 PRIME, Gaithersburg, USA) and the bead-milling TissueLyser system (Qiagen, Hilden, Germany). QuantiTect® Reverse Transcription Kit (Qiagen, Hilden, Germany) was used to generate cDNA. The primers (Microsynth, Balgach, Switzerland) used for the quantitative polymerase chain reaction (qPCR) are summarized in **Supplementary Table 1**. Quantitation of *CNR2, Iba1, TNF-α, MMP9, GFAP* and *MAP-2* mRNA expression was performed with the DyNAmo™ Flash SYBR® Green qPCR Kit (Thermo Scientific, Runcorn, UK) using a 7900 HT Fast Real-Time PCR System (Applied Biosystems, Carlsbad, USA). The amplification signals were detected in real-time, which permitted accurate quantification of the amounts of the initial RNA template during 40 cycles according to the manufacturer’s protocol. All reactions were performed in duplicates and in two independent runs. Quantitative analysis was performed using the SDS Software (v2.4) and a previously described 2−ΔΔCt quantification method [69]. The specificity of the PCR products of each run was determined and verified with the SDS dissociation curve analysis feature.

### *In vivo* MRI

Data were acquired at 24 h after reperfusion on a 7 T Bruker Pharmascan (Bruker BioSpin GmbH, Germany), equipped with a volume resonator operating in quadrature mode for excitation and a four element phased-array surface coil for signal reception and operated by Paravision 6.0 (Bruker BioSpin) [67, 70–72]. Mice were anesthetized with an initial dose of 4 % isoflurane in oxygen/air (200:800 ml/min) and maintained at 1.5 % isoflurane in oxygen/air (100:400 ml/min). Body temperature was monitored with a rectal temperature probe (MLT415, ADInstruments) and kept at 36.5 °C ± 0.5 °C using a warm water circuit integrated into the animal support (Bruker BioSpin GmbH, Germany). T_2_-weighted MR images were obtained using a spin echo sequence (TurboRARE) with an echo time 3 ms, repetition time 6 ms, 100 averages, slice thickness 1 mm, field-of-view 2.56 cm × 1.28 cm, matrix size 256 × 128, giving an in-plane resolution of 100 μm × 100 μm. For DWI, a four-shot spin echo–echo planar imaging sequence with an echo time = 28 ms, repetition time = 3000 [70, 71] acquired with a field-of-view of 3.3 cm × 2 cm and a matrix size of 128 × 128, resulting in a nominal voxel size of 258 μm × 156 μm. Diffusion-encoding was applied in the x-, y-, and z-directions with b-values of 100, 200, 400, 600, 800, and 1000 s/mm^2^, respectively, acquisition time 3 min 48 s. The ischemic lesion was determined as an area of significant reduction of the apparent diffusion coefficient (ADC) value compared with the unaffected contralateral side [73]. On T_2_-weighted images, the lesion was determined as an area of hyperintensities compared with the contralateral side.

### *In vivo* microPET studies

MicroPET/CT scans were performed at 24 h after reperfusion with a calibrated SuperArgus μPET/CT scanner (Sedecal, Madrid, Spain) with an axial field-of-view of 4.8 cm and a spatial resolution of 1.6–1.7 mm (full width at half maximum). tMCAO and the sham-operated C57BL/6J mice were anesthetized with ca. 2.5 % isoflurane in oxygen/air (1:1) during tracer injection and the whole scan time period. The formulated radioligand solution ([^18^F]FDG: 9.9-11 MBq or [^18^F]RoSMA-18-d6: 7.2-13 MBq) was administered via tail vein injection, and mice were dynamically scanned for 60 min. For blocking experiments, 1.5 mg/kg GW405833 was dissolved in a vehicle of 2 % Cremophor (v/v), 10 % ethanol (v/v), and 88 % water for injection (v/v) and injected together with [^18^F]RoSMA-18-d6. Body temperature was monitored by a rectal probe and kept at 37 °C by a heated air stream (37 °C). The anesthesia depth was measured by the respiratory frequency (SA Instruments, Inc., Stony Brook, USA). μPET acquisitions were combined with CT for anatomical orientation and attenuation correction. The obtained data were reconstructed in user-defined time frames with a voxel size of 0.3875 × 0.3875 × 0.775 mm^3^ as previously described [74].

### Triphenyltetrazolium chloride (TTC) staining

To assess the ischemic lesion severity in the brain of tMCAO mice and to validate the absence of lesion in the sham-operated mice, staining with TTC staining was performed. After measurements mice were euthanized, their brains were removed and 1-mm thick brain slices were obtained with a brain matrix. Slices were incubated in a 2.5 % TTC solution (Sigma-Aldrich, Switzerland) in PBS at 37 °C for 3 min. Photographs of the brain sections were taken. Edema-corrected lesion volumes were quantified as described [75].

### Biodistribution studies in the mouse brain

After PET/CT scanning of tMCAO mice at 24 h after reperfusion with [^18^F]RoSMA-18-d6, animals were sacrificed at 70 min post injection by decapitation. The spleen and brain regions of ischemic ipsilateral area and contralateral hemisphere were collected for analysis with a gamma counter. The accumulated radioactivities in the different tissues were expressed as percent normalized injected dose per gram of tissue normalized to 20 g body weight of the animals (norm. percentage injected dose per gram tissue (% ID/g tissue)).

### Data analysis and Statistics

Images were processed and analyzed using PMOD 4.2 software (PMOD Technologies Ltd., Zurich, Switzerland). The time−activity curves were deduced from specific volume-of-interest that were defined based on a mouse MRI T_2_-weighted image template [76]. Radioactivity is presented as standardized uptake value (SUV) (decay-corrected radioactivity per cm^3^ divided by the injected dose per gram body weight). [^18^F]RoSMA-18-d6 SUVR was calculated by using the midbrain in the corresponding hemisphere as reference brain region. For [^18^F]FDG PET, regional SUV was calculated. Two-way ANOVA with Sidak post-hoc analysis was used for comparison between groups (Graphpad Prism 9.0, CA, U.S.A).

## Results

### Increased expression of inflammation makers and neuronal damage after focal cerebral ischemia in tMCAO mice

mRNA levels were measured to address the question whether mouse non-ischemic and ischemic hemispheres differ in their expression levels of *CNR2* and other inflammatory genes. *CNR2* mRNA expression was increased to around 1.3 fold after 24 h reperfusion and at 48 h in the ipsilateral comparing to contralateral hemisphere (**Fig. 1a**). Similar 1.5-2.5 fold increases were observed in the mRNA expression of inflammatory markers including *TNF-α*, *Iba1, MMP9* and *GFAP* at 24 h and 48 h after reperfusion in the ipsilateral compared to contralateral brain region **(Fig. 1b–e**). *MAP-2* expression has been shown to be a reliable marker of neurons that undergo irreversible cell death [77, 78]. The neuron-specific *MAP-2* expression was markedly reduced in the ipsilateral compared to contralateral hemisphere at 24 h and 48 h after reperfusion (**Fig. 1f**). As similar *CNR2* mRNA expression were observed in 24 h and 48 h, our studies were performed at early time point of 24 h after reperfusion for investigating the functional, structural and molecular changes in the following experiments.

**Fig 1.**
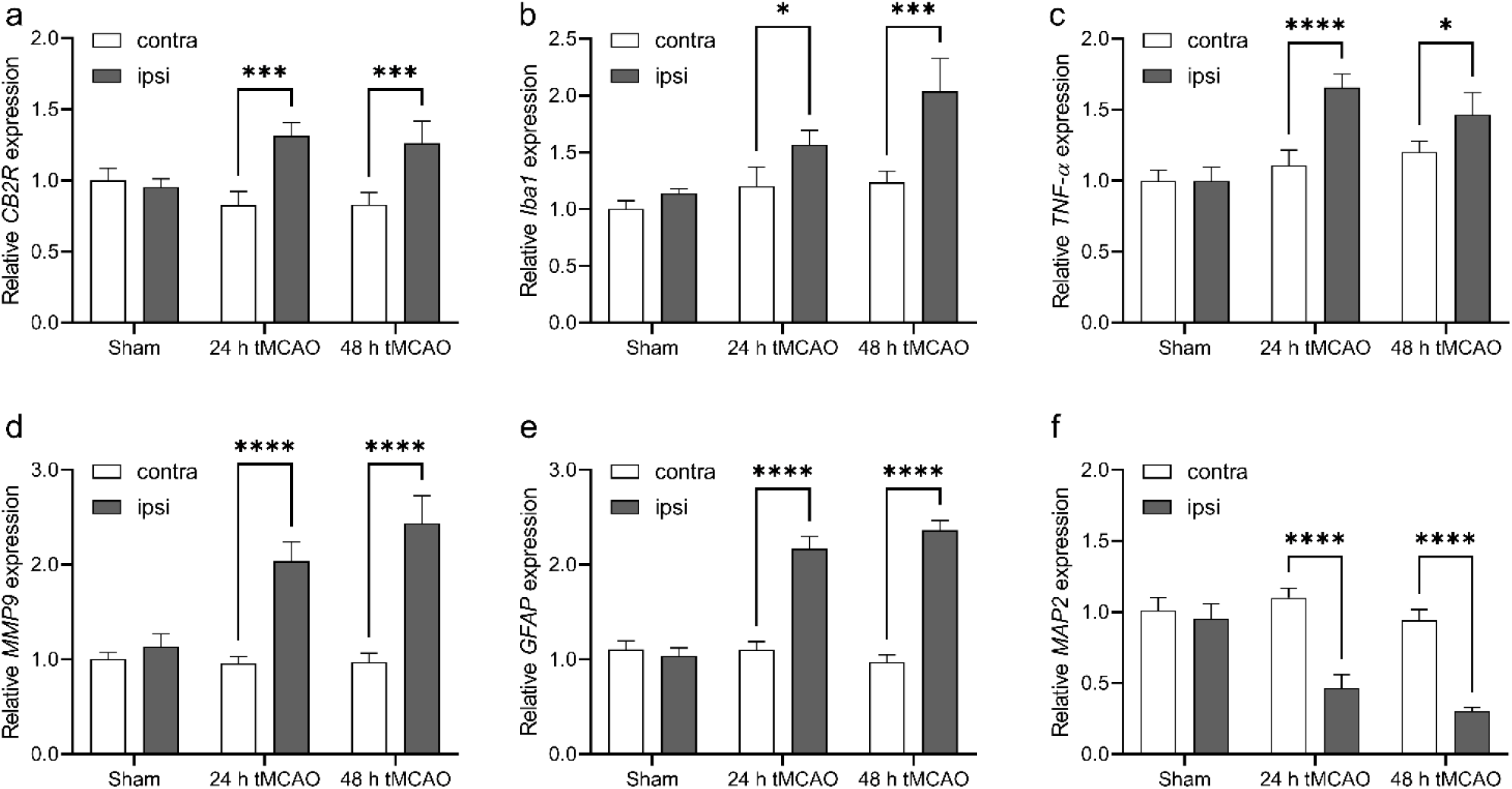
Relative mRNA levels of inflammatory markers and neuronal damage in sham-operated and tMCAO mouse brain in contra-and ipsilateral brain hemisphere at 24 h and 48 h after reperfusion. (a) *CNR2, (b) Iba1, (c) TNF-α, (d) MMP9, (e) GFAP* and (f) *MAP-2*. Values represent mean ± standard deviation. Expression levels were quantified by qPCR relative to β-actin.

### Reduced cerebral glucose metabolism and structural MRI lesion following tMCAO

Reduced [^18^F]FDG uptake was observed in the presumed MCA territory of the ipsilateral hemisphere in tMCAO mice, while there was no difference in [^18^F]FDG uptake between hemispheres in sham-operated mice (**Fig. 2a**). SUVs were significantly lower in the ipsilateral in the striatum in tMCAO compared to the contralateral side and compared to the same region in sham-operated mice 1.8 vs 1.4 (**Fig. 2b**). There were no differences in [^18^F]FDG uptake in the cortex and cerebellum between the ipsilateral and contralateral hemisphere in tMCAO mice and sham-operated mice. T_2_-weighted MRI and DWI imaging were performed in tMCAO and sham-operated animals at 24 h after reperfusion (**Fig. 2c**). The lesions in the ipsilateral side in the striatum and cortex were visible as areas of decreased values on the ADC maps calculated from DWI, and as areas of increased intensities on the T_2_-weighted MR images at 24 h after reperfusion following 1 h tMCAO (**Figs. 2c–d**). Ischemic lesions in the tMCAO were also seen *ex vivo* as white areas while viable tissue appeared red in TTC stained brain sections(**Fig. 2e**). Homogenous deep red color was observed across both hemispheres in sham-operated mice, verifying the absence of any lesion. The hemispheric lesion volumes in tMCAO mice were 42.8 ± 10.2 % (mean ± standard deviation).

**Fig 2.**
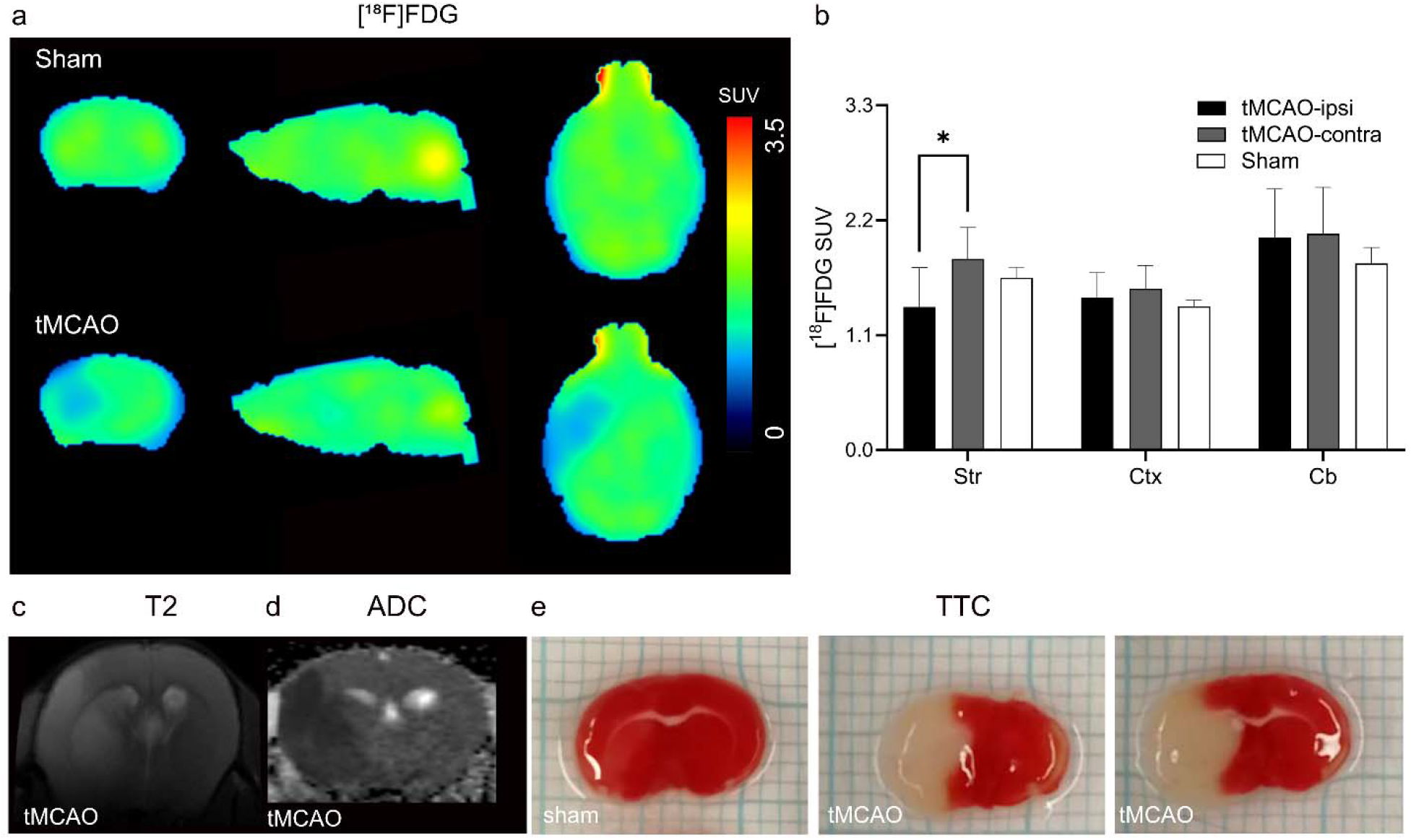
*In vivo* MRI and [^18^F]FDG PET in tMCAO mouse brain. (**a**) Representative PET images of coronal, sagittal and horizontal mouse brain sections after intravenous injection of [^18^F]FDG in sham-operated and tMCAO mice. The radiosignals were averaged from 21-61 min; (**b**) [^18^F]FDG accumulation (SUV) at different mouse brain regions (Str: striatum; Ctx: cortex; Cb: cerebellum) in sham and tMCAO mice. Significantly reduced [^18^F]FDG accumulation was observed in the ipsilateral striatum compared to contralateral side in tMCAO mice; (**c-e**) *In vivo* T_2_-weighted image, ADC map and *ex vivo* TTC stained brain sections, indicating the delineation in tMCAO mice. TTC: 2,3,5-triphenyltetrazolium chloride; ADC: apparent diffusion coefficient; SUV: standard uptake value.

### Increased [^18^F]RoSMA-18-d6 retention in the striatum after tMCAO

To analyze the distribution of [^18^F]RoSMA-18-d6 in tMCAO mice brain, dynamic μPET/CT scans were performed at 24 h after reperfusion. The standard uptake values (SUVs) of [^18^F]RoSMA-18-d6 did not reveal significant difference in various brain regions of tMCAO mice (**Supplementary Fig 1**). However, we found a reduced uptake at early time frame (1-3 min) and a similar uptake after 7 min in the ipsilateral side compared to that of contralateral side (Fig. 3a). Thus, to exclude the perfusion influence, we averaged the brain signals from 21-61 min and selected the midbrain as the reference region. Higher [^18^F]RoSMA-18-d6 SUVR was observed in the ischemic ipsilateral striatum compared to the contralateral striatum (two-way ANOVA with Sidak multiple comparison correction, 0.97± 0.02 vs 0.87 ± 0.06, p = 0.0274), but not in other brain regions such as cortex (**Fig. 3b, c**). The increased signals at ischemic ipsilateral striatum, however, could not be blocked by the selective CB_2_R agonist GW405833 (**Fig. 3c**).

**Fig 3.**
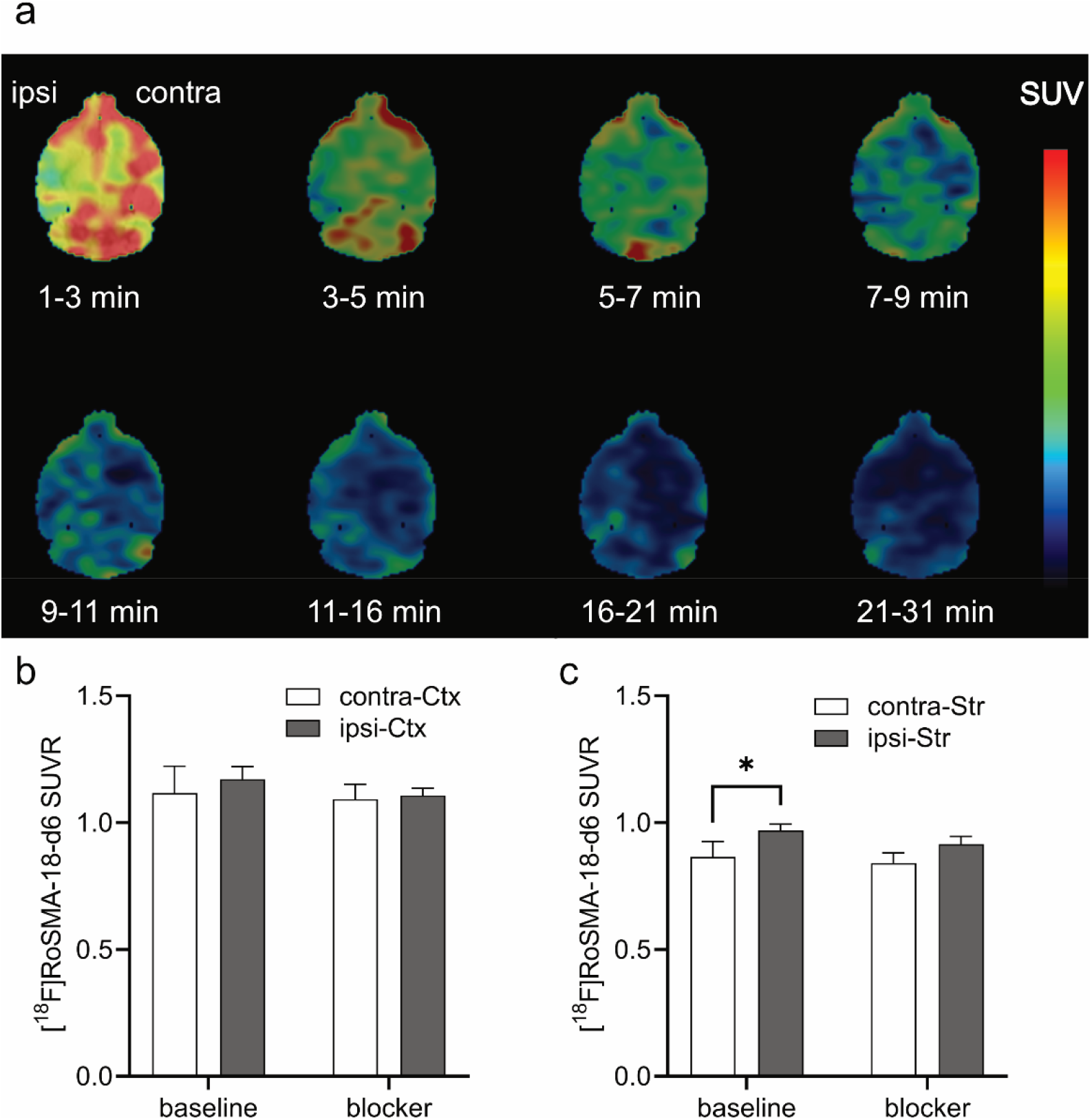
*In vivo* microPET imaging of tMCAO mouse brain using [^18^F]RoSMA-18-d6. (**a**) Representative PET images of horizontal mouse brain sections at different time frames after intravenous injection of [^18^F]RoSMA-18-d6; SUV: 0-0.5; (**b, c**) Ratios of [^18^F]RoSMA-18-d6 uptake under baseline and blockade conditions in cortex and striatum. Significantly higher [^18^F]RoSMA-18-d6 standard uptake value ratio (SUVR) was observed in the ischemic ipsilateral striatum under baseline conditions, but not in the ipsilateral cortex. Midbrain was used as reference brain region for SUVR calculation.

At the end of the *in vivo* experiments, we dissected the mice to verify the activity accumulation and specificity of [^18^F]RoSMA-18-d6 in the spleen and different brain regions with a gamma counter. In line with the results obtained from the averaged SUVRs in the tMCAO mouse brain, the radioactivity in the ipsilateral side was indeed significant higher than that of the contralateral hemisphere (0.037 ± 0.007 vs 0.026 ± 0.003, n = 5 each group), but no blockade effect was seen under blocking conditions (**Fig. 4a**). As expected, radioactivity in the CB_2_R-rich spleen was much higher than the brain and 58 % of the signals was blocked by co-injection of CB_2_R specific ligand GW405833, demonstrating specific target engagement of [^18^F]RoSMA-18-d6 *in vivo* (**Fig. 4b**).

**Fig 4.**
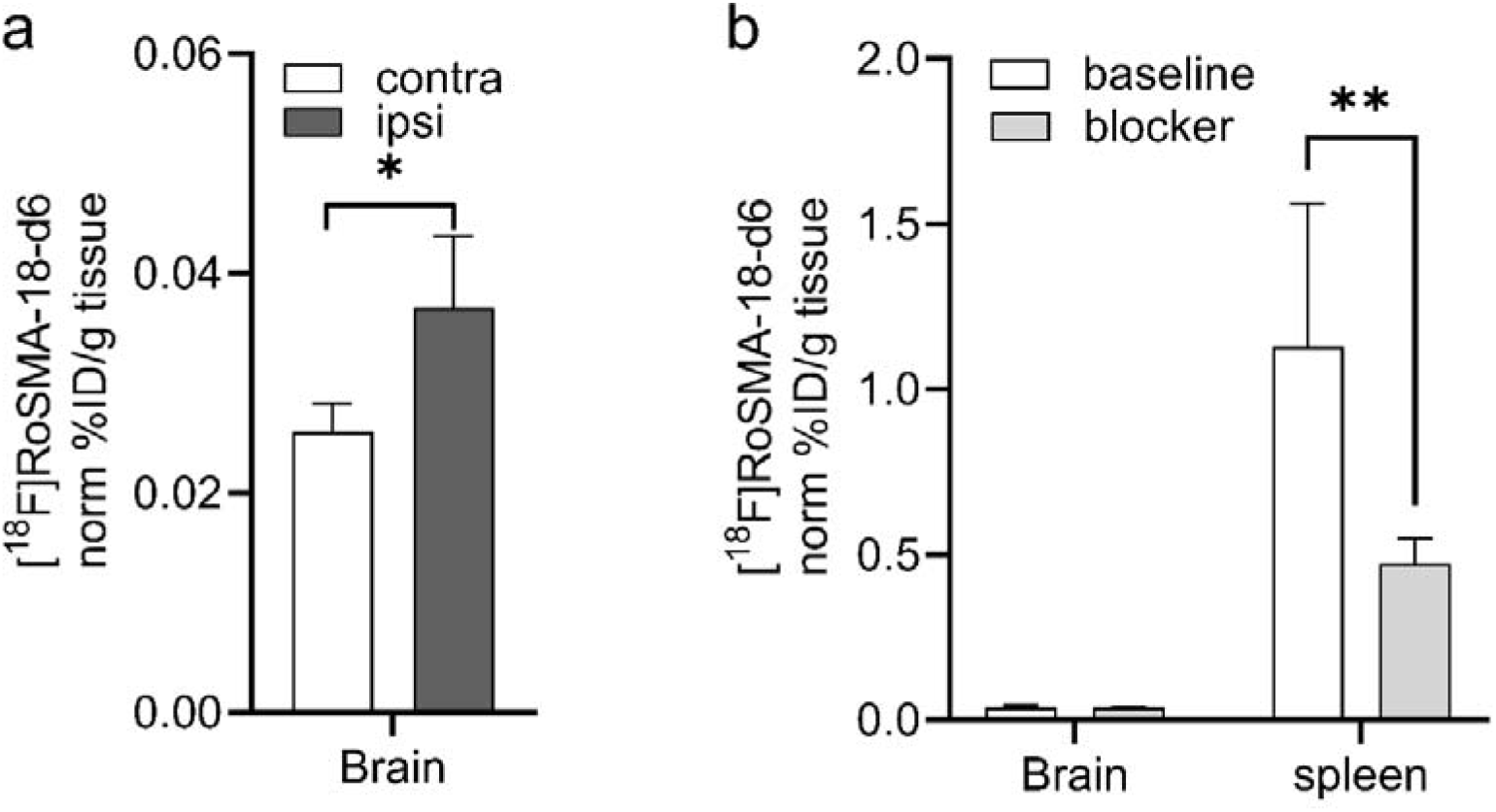
*Ex-vivo* biodistribution of [^18^F]RoSMA-18-d6 in the brain and spleen of tMCAO mouse. Animals (n=4) were sacrificed at 70 min post-injection, the spleen and brain regions were dissected and analyzed with a gamma counter. (**a**) Higher [^18^F]RoSMA-18-d6 binding (norm% ID/g tissue) was detected in the ipsilateral vs contralateral hemisphere under baseline conditions. (**b**) In the spleen about 58 % of the [^18^F]RoSMA-18-d6 binding (norm% ID/g tissue) was blocked. No significant blocking was observed in the brain. Data are presented as the mean of the percentage of injected dose per gram tissue normalized to 20 g body weight; mean ± standard deviation. % ID/g: percentage injected dose per gram.

## Discussion

This study assessed the utility of CB_2_R PET tracer [^18^F]RoSMA-18-d6 for imaging tMCAO mouse at subacute stage, concomitant with decreased CMRglc levels and formation of a structural lesion. Previous PET imaging of stroke animal models led to inconclusive results. In a rat model of photothrombotic stroke at 24 h after surgery, increased [^11^C]NE40 (CB_2_R tracer) uptake and unvaried [^11^C]PK11195 (TSPO tracer) uptake were reported [79]. In another study, [^11^C]NE40 uptake did not show any difference in the same rat model of photothrombotic stroke [80]. Moreover, reduced [^11^C]A836339 (CB_2_R tracer) uptake was reported in a focal tMCAO rat model over 1-28 days after occlusion [51]. Possible reasons for these different observations include the time point of assessment, different methods for inducing acute stroke (transient or permanent ischemia) resulting in variations of ischemic severity and levels of inflammatory-cell expression [43].

CB_2_R has negligible expression in the mouse brain and is mainly expressed in the spleen under physiological conditions [30, 36, 60–65, 81]. Under neuroinflammatory conditions, CB_2_R is upregulated in activated microglial cells. In this study, we used quantitative real-time polymerase chain reaction to measure gene expression levels of *CNR2, TNF-α, Iba1, MMP9, GFAP* and *MAP-2* at 24 h and 48 h. All tested inflammatory markers displayed increased mRNA levels in the ipsilateral brain hemisphere, in agreement with the reported findings in tMCAO mouse model [29, 45, 82, 83]. In line with the increased *CNR2* gene expression levels, significantly higher [^18^F]RoSMA-18-d6 SUVR (standard uptake value ratio) was observed in striatum at ipsilateral vs contralateral under baseline conditions in our PET studies. The 50 % reduction of the neuronal marker *MAP-2* indicated neuronal damage.

The dynamic μPET scan using [^18^F]RoSMA-18-d6 indicated a reduced perfusion in the lesion brain regions at the first time frame of 1-3 minutes. This is probably due to the changes of microvascular response (no-reflow phenomenon) and the reduction in neuronal activity. Taking the midbrain as the reference region, the ratios of SUV averaged from 21-61 min revealed increased [^18^F]RoSMA-18-d6 SUVR in the ipsilateral ischemic striatum compared to that of the contralateral side. Our *ex vivo* bio-distribution studies confirmed the difference of the radioactivity distribution in the left and right brain hemisphere. The *in vivo* specificity of [^18^F]RoSMA-18-d6 towards CB_2_R is evidenced by a 58 % reduction in radioactivity in the mouse spleen under blockade conditions in *ex vivo* biodistribution studies. Underlying reasons for the lack of specificity of [^18^F]RoSMA-18-d6 in the mouse brain may because 1) the increased tracer availability in the blood induced by blocking the CB_2_R peripheral targets in the presence of the blocker GW405833; and 2) the relatively low brain uptake of our CB_2_R-selective radioligand [^18^F]RoSMA-18-d6 in the mouse brain resulted in undetectable changes of radiosignals under baseline and blockade conditions. Notably, the time-activity curves of [^18^F]RoSMA-18-d6 in tMCAO mouse brain showed remarkably higher initial brain uptake under blockade conditions than the baseline in both sides of the mouse brain (**Supplementary Fig 1**), indicating the influence of blocking CB_2_R target in the peripheral organs on the availability of radiotracer concentrations in the blood. In our previous studies with Wistar rat, the spleen uptake of [^18^F]RoSMA-18-d6 was blocked by nearly 90 % suggesting a high possibility of species difference of [^18^F]RoSMA-18-d6 binding [57]. Therefore, we speculate that rat stroke models might be superior to mice models for imaging neuroinflammation with CB_2_R PET tracers.

We observed that [^18^F]FDG measure of CMRglc was reduced in the ischemic areas i.e. ipsilateral striatum of the tMCAO mice at 24 h after reperfusion. The reduced CMRglc was reported in many earlier studies in disease animal models and in stroke patients [84–87], masking CMRglc reduction of neuronal tissue in the brain At an extended time points of the recovery stage from day 4 - 40, an increased CMRglc level was reported in the ischemic regions due to the increased consumption from inflammatory cells along with microglial activation [88–90].

There are several limitations in the current study. 1) As there is no reliable specific CB_2_R antibody, we did not include immunohistochemical staining for CB_2_R protein distribution in the mouse brain. The qPCR measures of *CNR2* mRNA level provided an alternative readout, but do not provide spatial distribution of cerebral CB_2_R expression. 2) Due to the logistic barrier, MRI and μPET/CT scans were performed with different cohorts of animals. Nevertheless, standard operating procedures for the surgery were used. 3) Our *in vivo* data with tMCAO mice were collected at 24 h after surgery, longitudinal imaging of tMCAO mice with [^18^F]RoSMA-18-d6 along with structural and functional readout will provide further insight into the spatial-temporal dynamics of CB_2_R expression in the brain.

## Conclusion

Our newly developed CB_2_R PET tracer [^18^F]RoSMA-18-d6 revealed limited utility to image neuroinflammation in the ischemic ipsilateral of the tMCAO mice at 24 h after reperfusion. Although lesion regions in tMCAO mouse brain could be followed by the ratios of averaged SUVs from 21-61 with midbrain as the reference region, the in-vivo specificity of [^18^F]RoSMA-18-d6 was confirmed only in the CB2R-rich spleen. Different neuroinflammatory animal models which has comparable neuronal numbers in the lesion regions are recommended for evaluation of CB_2_R in further PET imaging studies.

## Acknowledgements

The authors acknowledge Yingfang He, Annette Krämer at Center for Radiopharmaceutical Sciences, Department of Chemistry and Applied Biosciences, ETH Zurich; Dr Mark Augath, at the Institute for Biomedical Engineering, ETH Zurich & University of Zurich for technical assistance.

## Abbreviations

ADC: Apparent diffusion coefficient
CB_2_R: Cannabinoid type 2 receptors
CBF: Cerebral blood flow
CMRglc: Cerebral metabolic rate of glucose
CT: computed tomography
DW: Diffusion weighted
FDG: fluorodeoxyglucose
% ID/g tissue: Injected dose per gram tissue
MRI: Magnetic resonance imaging
PET: Positron emission tomography
SUV: Standardized uptake value
SUVR: Standard uptake value ratio
tMCAO: transient middle cerebral artery occlusion
TSPO: Translocator protein
TTC: Triphenyltetrazolium chloride

## Availability of data and materials

The data used and analyzed in the current study are available from the corresponding authors upon request.

## Ethics approval and consent to participate

All experiments were performed in accordance with the Swiss Federal Act on Animal Protection and were approved by the Cantonal Veterinary Office Zurich (permit number: ZH018/14 and ZH264/16).

## Consent for publication

Not applicable.

## Authors’ contribution

RN, JK, LM, SMA designed the study; RN, KC, AH, AMH, GL, LM performed the experiment, RN, LM performed data analysis, RN, JK, LM wrote the initial manuscript. All authors read and approved the final manuscript.

## Additional files

Additional file 1: Supplementary Figure 1. Time activity curves of [^18^F]RoSMA-18-d6 *in vivo* microPET imaging of tMCAO mouse brain. (**a-d**) In the cortex, striatum, cerebellum and midbrain under baseline and blockade conditions. No difference in [^18^F]RoSMA-18-d6 SUV was observed in different brain regions at ipsilateral vs contralateral side under baseline or blockade conditions. Data represent mean ± standard deviation.

Additional file 2: Supplementary Table 1. Primers used for the quantitative polymerase chain reaction assay on mouse brain tissue

