## Supplementary files for "*In vivo* imaging of cannabinoid type 2 receptors, functional and structural alterations in mouse model of cerebral ischemia by PET and MRI"

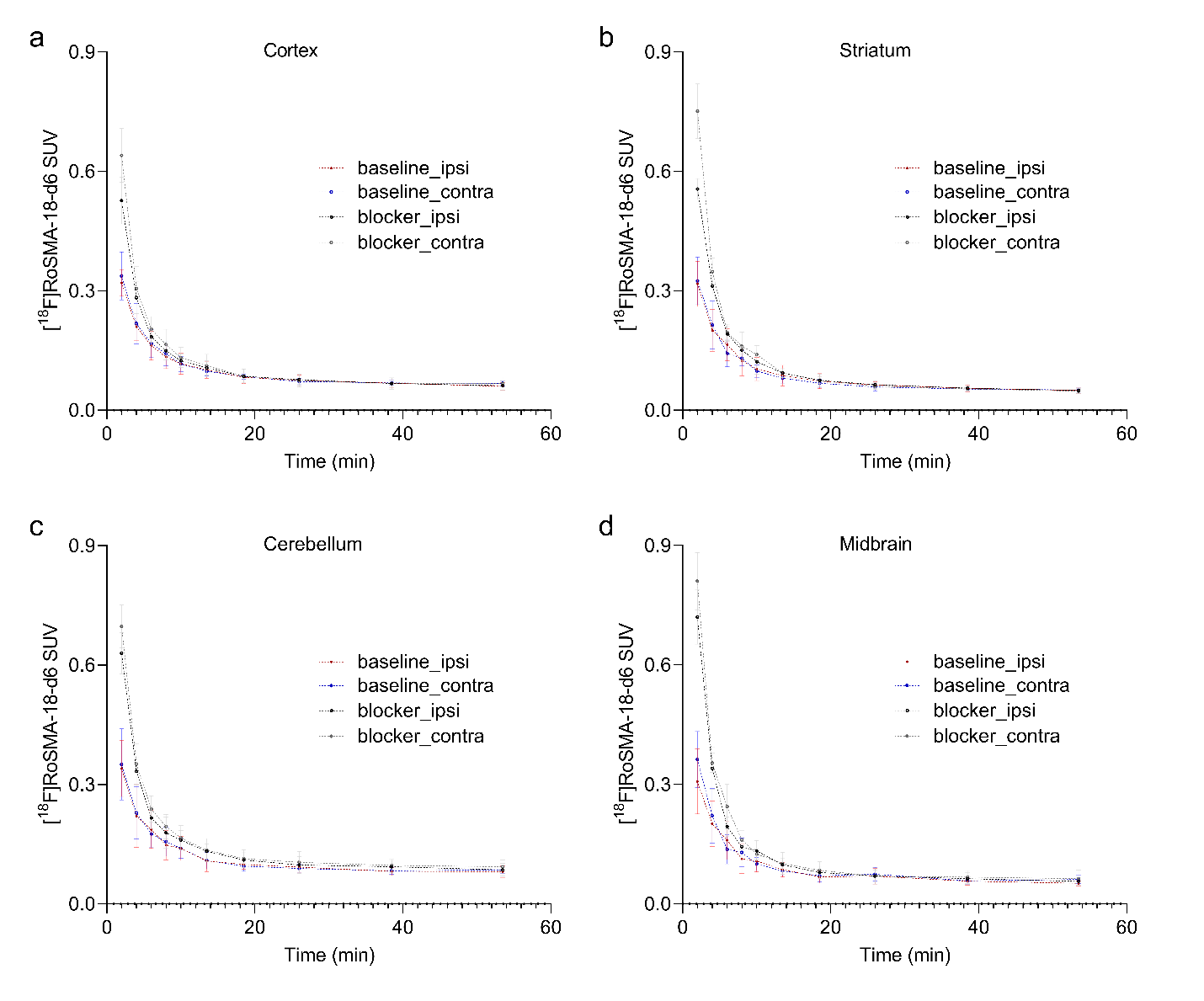

**Supplementary Figure 1.** Time activity curves of [^18^F]RoSMA-18-d6 *in vivo* microPET imaging in tMCAO mouse brain. (**a-d**) In the cortex, striatum, cerebellum and midbrain under baseline (n = 6) and blockade (n = 4) conditions. No difference in [^18^F]RoSMA-18-d6 SUV was observed in different brain regions at ipsilateral vs contralateral side under baseline or blockade conditions. Data represent mean ± standard deviation.

**Supplementary Table 1: Primers used for the quantitative polymerase chain reaction assay on mouse brain tissue**

| RNA | Primer |
| --- | --- |
| *beta-actin (ACTB)* | forward 5′-AGACCTCTATGCCAACACAGT-3′, reverse 5′-TGCTAGGAGCCAGAGCAGTAA-3′ |
| *CNR2 (CB2R)* | forward 5′-CTACAAAGCTCTAGTCACCCGT-3′, reverse 5′-CCATGAGCGGCAGGTAAGAAA-3’ |
| *ionized calcium binding adaptor molecule 1 (Iba1)* | forward 5’-GTCCTTGAAGCGAATGCTGG-3’, reverse 5’-CATTCTCAAGATGGCAGATC-3’ |
| *Tumor necrosis factor (TNF-α)* | forward 5′-AATGGCCTCCCTCTCATCAGTT-3′, reverse 5′-CCACTTGGTGGTTTGCTACGA-3′ |
| *Matrix metallopeptidase 9 (MMP9)* | forward 5’-AACATCTGGCACTCCACACC-3’, reverse 5’-GCAGAAGTTCTTTGGCCTGC-3’ |
| *Glial fibrillary acidic protein (GFAP)* | forward 5’-CGGAGACGCATCACCTCTG-3’, reverse 5’-TGGAGGAGTCATTCGAGACAA-3’ |
| *microtubule-associated protein 2 (MAP-2)* | forward 5’-GCCAGCCTCAGAACAAACAG-3’, reverse 5’-AAGGTCTTGGGAGGGAAGAAC-3’ |
